## Supplementary material for "Redefining hypo- and hyper-responding phenotypes of CFTR mutants for understanding and therapy": Suppl. Data

### Figure S1

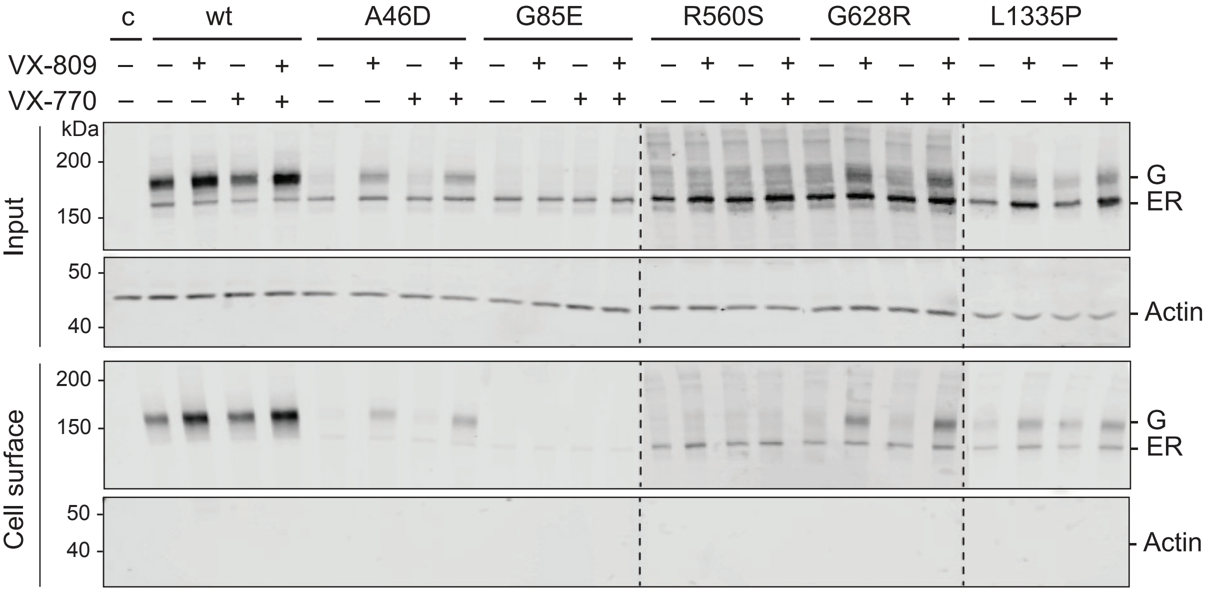
c
wt
A46D
G85E
R560S
G628R
L1335P
VX-809
VX-770
kDa
200
150
50
40
Input
Cell surface
G
ER
Actin

### Figure S2

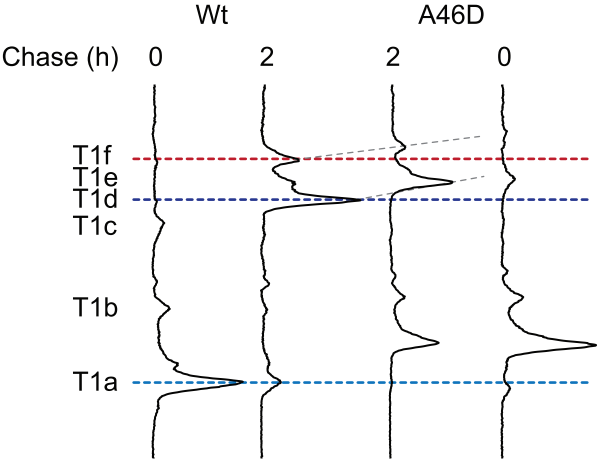

### Figure S3

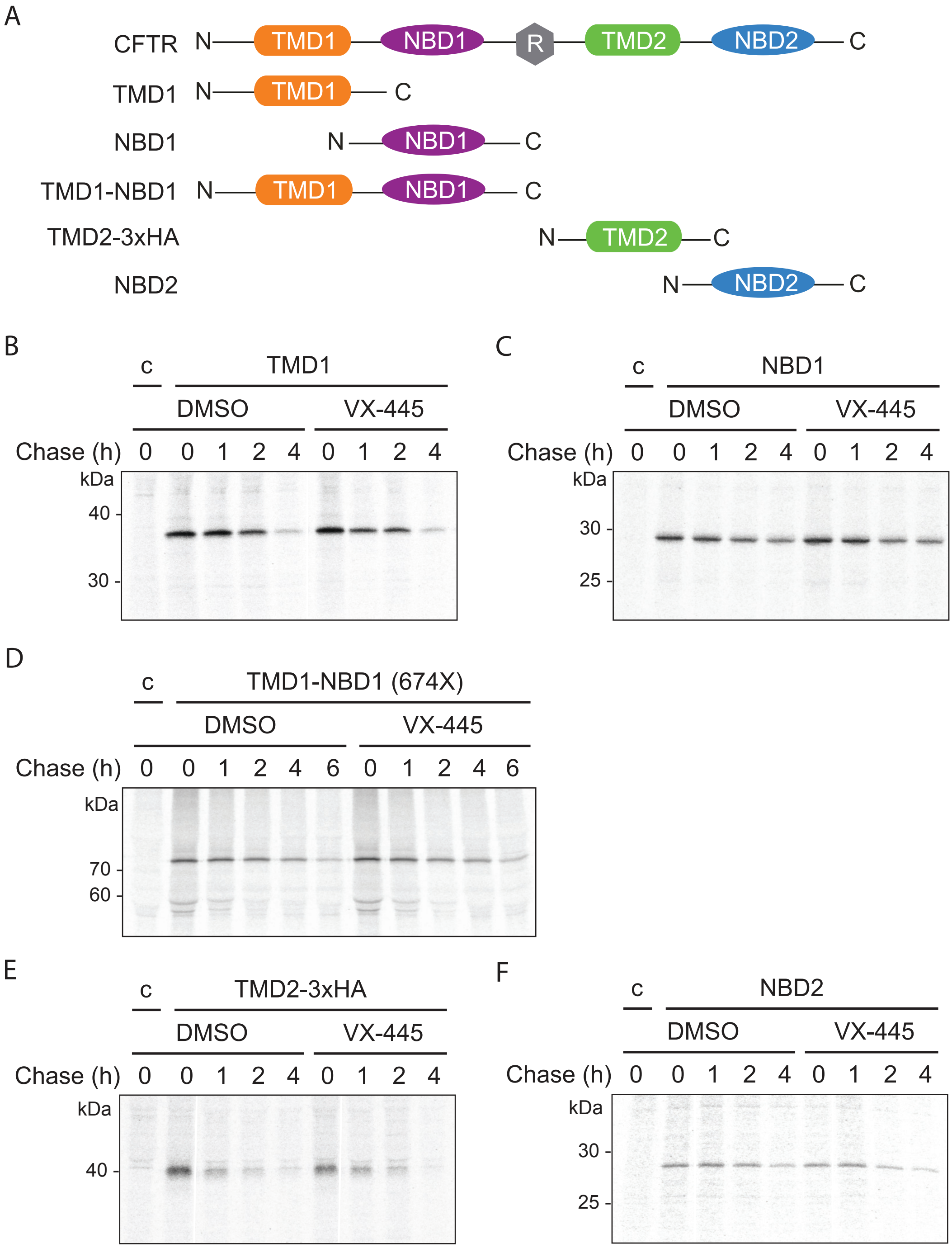

A
CFTR N TMD1 NBD1 R TMD2 NBD2 C
TMD1 N TMD1 C
NBD1 N NBD1 C
TMD1-NBD1 N TMD1 NBD1 C
TMD2-3xHA N TMD2 C
NBD2 N NBD2 C
B
c TMD1
DMSO VX-445
Chase (h) 0 0 1 2 4 0 1 2 4
kDa
40
30
C
c NBD1
DMSO VX-445
Chase (h) 0 0 1 2 4 0 1 2 4
kDa
30
25
D
c TMD1-NBD1 (674X)
DMSO VX-445
Chase (h) 0 0 1 2 4 6 0 1 2 4 6
kDa
70
60
E
c TMD2-3xHA
DMSO VX-445
Chase (h) 0 0 1 2 4 0 1 2 4
kDa
40
F
c NBD2
DMSO VX-445
Chase (h) 0 0 1 2 4 0 1 2 4
kDa
30
25

**Supplementary figure 1: Figure 1 with darker exposure for R560S and G628R lanes**
**(A)** Figure 1B, but with decreased brightness for R560S and G628R CFTR panels.

**Supplementary figure 2: Lane profiles belonging to Figures 2 and 4**
TMD1 fragments from A46D at 0-h and 2-h chase times deviated from the typical wild-type-like pattern. Lane profiles from Figure 4B, C without modulators show the upshift of the fragments that contain the A46D mutation. T1a does not contain A46D and disappears because L49 becomes shielded in the mutant. Red dotted line indicates mobility of wt T1f; dark blue dotted line indicates mobility of wt T1d, light blue dotted line indicates mobility of wt T1a, dark grey dotted lines represent shifts in T1f and T1d.

**Supplementary figure 3: Single and multi-domain CFTR constructs are not stabilized in the presence of VX-445**
**(A)** Schematic representation of constructs used in (B-F). **(B)** HEK293T cells expressing isolated TMD1 were radiolabeled for 5 minutes and chased for 0, 1, 2, or 4 hours. VX-445 (3 µM) was added during starvation, pulse, and chase. Cycloheximide (1 mM) was added during the chase only. CFTR was immunoprecipitated from detergent cell lysates using E1-22 and analyzed on 12% SDS-PA gels. **(C-F)** As in (B), intracellular stability of **(C)** Wild-type isolated NBD1, immunoprecipitated with MrPink, **(D)** TMD1-NBD1 (674X), also chased for 6 h and immunoprecipitated with MrPink, **(E)** Isolated TMD2-3xHA CFTR, immunoprecipitated with TMD2C, and **(F)** Isolated NBD2, immunoprecipitated with 596.

**Table S1. Primers used for cloning of the indicated CFTR constructs**

F, forward primer; R, reverse primer.

| CFTR construct | Primer sequence 5' to 3' |
|---|---|
| A46D | F: GTTGATTCTGATGACAATCTATCTGAA |
| | R: TTCAGATAGATTGTCATCAGAATCAAC |
| G461R | F: TCCACTAGAGCAGGCAAGA |
| | R: TCTTGCCTGCTCTAGTGGA |
| F508del | F: CCATTAAAGAAAATATCATTGGTGTTTCCTATGATGAATATAG |
| | R: CTATATTCATCATAGGAAACACCAATGATATTTTCTTTAATGG |
| R560S | F: TCTTTAGCAAGCGCAGTATACAAAGATG |
| | R: CATCTTTGTATACTGCGCTTGCTAAAGA |
| G628R | F: AGCAGCTATTTTTATAGGACATTTTCAG |
| | R: CTGAAAATGTCCTATAAAAATAGCTGCT |
| G1249R | F: ACTGGATCAAGGAAGAGTACTTTG |
| | R: CAAAGTACTCTTCCTTGATCCAGT |
| L1335P | F: TTTCCTGGGAAGCCTGACTTTGT |
| | R: ACAAAGTCAGGCTTCCCAGGAAA |
| NBD2-CFTR (1202-1480) | F: GGGGCGGCCGCGACCATGGACATCTGGCCCTCAGGGGGCCAAATG |
| | R: CCTCTGGAGATATCGTCGACAAGCTTATCGATGCG |
